# *In- & Out-Cloning*: Plasmid toolboxes for scarless transcription unit and modular Golden Gate acceptor plasmid assembly

**DOI:** 10.1101/2024.06.22.600171

**Authors:** Stijn T. de Vries, Tania S. Köbel, Ahmet Sanal, Daniel Schindler

**Affiliations:** Max Planck Institute for Terrestrial Microbiology, Karl-von-Frisch-Str. 10, 35043 Marburg, Germany; Center for Synthetic Microbiology, Philipps-University Marburg, Karl-von-Frisch-Str. 14, 35032 Marburg, Germany

**Keywords:** DNA assembly, Golden Gate cloning, SapI, Modular Cloning, Synthetic biology

## Abstract

Golden Gate cloning has become one of the most important DNA assembly strategies. The construction of standardized and reusable part libraries, their assembly into transcription units, and the subsequent assembly of multigene constructs is highly reliable and sustainable. Researchers can quickly construct derivatives of their assemblies or entire pathways, and importantly, the standardization of Golden Gate assemblies is compatible with laboratory automation. Most Golden Gate strategies rely on four nucleotide overhangs generated by commonly used Type IIS enzymes. However, reduction to three nucleotide overhangs allows the use of codons as fusion sites and reduces potential scar sequences. This is particularly important when studying biological functions, as additional nucleotides may alter the structure or stability of the transcribed RNA. To address this issue we use SapI, a Type IIS enzyme generating three nucleotide overhangs, for transcription unit assembly, allowing for codon-based fusion in coding sequences. We created a corresponding plasmid toolbox for basic part generation and transcription unit assembly, a workflow we term *In-Cloning. In-Cloning* is downstream compatible with the Modular Cloning standard developed by Sylvestre Marillonnet’s group for standardized assembly of multigene constructs. However, the multigene construct plasmids may not be compatible for use with the model organism of choice. Therefore, we have developed a workflow called *Out-Cloning* to rapidly generate Golden Gate acceptor plasmids. *Out-Cloning* uses standardized plasmid parts that are assembled into Golden Gate acceptor plasmids using flexible linkers. This allows the systematic construction of acceptor plasmids needed to transfer assembled DNA into the organism of interest.

## Introduction

One of the core elements of synthetic biology is the efficient and sustainable assembly of standardized DNA constructs. Golden Gate cloning, a DNA assembly method based on Type IIS enzymes, has become one of the most important techniques for efficient and sustainable DNA assembly (1). The advantage is its hierarchical and modular nature, which allows to divide a transcription unit (TU) into standardized positions, e.g. promoter, coding sequence (CDS) and terminator. For each position, reusable part libraries can be generated, allowing flexible, on-demand construction of TUs to efficiently address research questions. This approach can dramatically reduce the cost of sequence validation, as only basic parts need to be sequence validated. Based on Golden Gate cloning, the Modular Cloning (MoClo) system has been developed that allows the assembly of multiple TUs into higher order constructs, theoretically of infinite size (2). Several Type IIS enzymes are used for Golden Gate cloning, most of which generate four nucleotide (nt) overhangs (fusion sites). SapI is currently not widely used, but due to its 3 nt fusion sites it allows for a seamless start and stop codon cloning approach (3). The reduced fusion sites allow for codon-based cloning and have advantages for construction where minimal scar sequences are important, for example, for the construction and application of synthetic small regulatory RNAs where a single alteration of the sequence can cause a drastic decrease in functionality (4). Recently, researchers have determined the fidelity of fusion sites, and their data allow researchers to select optimal fusion sites to achieve maximum fidelity for Golden Gate cloning (5-7).

Here we present our plasmid toolbox for Golden Gate cloning using 3 nt fusion sites for the generation of TUs (“*In-Cloning*”) and the complementary toolbox for on demand generation of acceptor plasmids (“*Out-Cloning*”). The plasmid set is compatible with the MoClo standard developed by the group of Sylvestre Marillonnet and used in our previous work, but can be adapted to any other standard if required (2,8). The MoClo concept allows the generation of large DNA assemblies after iterative rounds of cloning. However, the assembly plasmids are not necessarily relevant for applications in the target organism and often specific acceptor plasmids are required. Therefore, *Out-Cloning* addresses this issue with its growing part library to create Golden Gate acceptor plasmids for use in the target organism on demand. The individual elements of the acceptor plasmids are positioned in the plasmid assembly using flexible linker, giving the user maximum freedom to generate acceptor plasmids. We routinely use *In-& Out-Cloning* for our research projects and describe our plasmid toolbox here to share this utility with the community.

## Methods

### Culture conditions, strains and plasmids used in this study

All yeast strains derived from *S. cerevisiae S288C* and were grown at 30 °C unless otherwise noted; all strains used and constructed in this study are listed in Table S1. Standard *E. coli* laboratory strains were used for propagation and archiving of plasmid DNA; all strains used and constructed in this study are listed in Table S1. Yeast cells were grown in either YEP/YPD media (10 g/L yeast extract, 20 g/L peptone, and with or without 0.64 g/L tryptophan) or Synthetic Complete (SC) media (1.7 g/L yeast nitrogen base without amino acids and without ammonium sulfate, 5 g/L ammonium sulfate, appropriate Synthetic Complete drop-out mixture (9); all components used were provided by Formedium) lacking the indicated amino acids, both supplemented with 2% glucose unless otherwise stated. Bacterial cultures were grown in LB medium (1% [w/v] bacto-tryptone, 1% [w/v] NaCl, 0.5% [w/v] yeast extract) at 37 °C with the indicated antibiotics (100 μg/mL ampicillin, 25 μg/mL chloramphenicol, 25 μg/mL gentamicin, 50 μg/mL kanamycin, 120 μg/mL spectinomycin). Bacterial and yeast liquid cultures were grown according to standard procedures in glass tubes in a custom-built roller drum (similar to New Brunswick TC-7) or in shaking incubators (Innova 42R, New Brunswick) at 200 rpm, unless otherwise stated. Standard solid media contained 2% agar. All plasmids used and generated in this study are listed in Table S2 and S3, respectively. A detailed description of the plasmids generated is provided in Supporting Data S1, and the sequences are provided as GenBank files in Supporting Data S2.

### Oligodeoxynucleotides and gene synthesis

All oligodeoxynucleotides were ordered in 25 or 100 nM scale as standard desalted oligonucleotides from Integrated DNA Technologies (IDT, Coralville, USA) with standard desalting purification. Only diagnostic and relevant oligonucleotides are provided in Table S4, oligonucleotides used for plasmid construction are not listed. However, all plasmid files are provided as GenBank files in Supporting Data S2. Gene synthesis was obtained as eBlocks or gBlocks from IDT (Coralville, Iowa, USA), or Gene Fragments from Twist Bioscience (South San Francisco, California, USA). Coding sequences were codon-matched and relevant Type IIS recognition sites (BbsI [BpiI], BsaI, Esp3I [BsmBI] and SapI) were excluded using DNA Chisel prior DNA synthesis (10).

### Extraction and purification of plasmid DNA and PCR products

Plasmid DNA and PCR product purification was performed by an open source magnetic bead purification procedures using carboxylated SeraMag Speed Beads (45152105050250 [cat#], Cytiva, Marlborough, USA) (11). Detailed, stepwise protocols are provided at http://www.bomb.bio. For plasmid extractions, cultures were grown in SMM medium (SMM: 16 g/L tryptone, 10 g/L yeast extract, 5 g/L glycerol, and 1× M9 salts) with the appropriate antibiotic in 2-3 mL in glass tubes on a standard shaker at 37 °C, 200 rpm, or in 1 mL in 96-well deep well plates (12). Deep well plates were inoculated overnight on an Infors HT shaker at 800 rpm, 3 mm shaking, and 80% humidity at 37 °C and sealed with a gas permeable seal (AB-0718, ThermoFisher Scientific, Waltham, USA). Plasmids and basic parts were verified using Sanger sequencing services and/or in-house Nanopore amplicon sequencing procedures (13,14). All generated plasmids are listed in Table S3, and their sequences are provided as annotated GenBank files in the Supporting Data S2.

### Gibson assembly based plasmid construction

Gibson assembly of plasmids was performed as previously described (12), and all resulting plasmid maps are provided in Supporting Data S2 as annotated GenBank files. Briefly, Gibson assembly was performed using an in-house prepared reaction mix in 10 or 20 μL total volume. All enzymes were provided by New England Biolabs (NEB, Ipswitch, USA), and the reaction mix and procedure were performed according to Gibson *et al*. (2009) (15). DNA fragments were amplified using Q5 DNA polymerase (NEB) or an in-house purified proofreading DNA polymerase. Gibson assemblies were transformed into in-house generated RbCl competent *E. coli* cells and selected on LB plates containing the appropriate antibiotic at 37 °C (12,16).

### Blunt-end ligation cloning for basic part generation

Level 0 parts can be constructed by blunt-end ligation in case of single fragment cloning. The primers for the PCR amplicons must contain the appropriate overhangs (*cf*. Figure 1 and Figure S2). The oligonucleotides or the resulting amplicons were phosphorylated with T4 polynucleotide kinase (PNK) (NEB) according to the manufacturer guidelines. The Level 0 plasmid was amplified with the primer pair SLo0765/SLo0766 followed by DpnI (NEB) digestion prior to ligation (any Level 0 plasmid will do, e.g. pSL099 because the fusion sites are omitted and provided by the part amplicon). PCR fragments were purified using the BOMB PCR clean-up procedure (11). The phosphorylated part amplicon and PCR amplified plasmid were ligated with T4 ligase (NEB) for 2 h at room temperature in a 4:1 ratio according to the manufacturer guidelines. Transformation was performed in in-house generated RbCl competent *E. coli* cells (12,16). Selection was performed overnight on LB plates at 37 °C with the respective antibiotic (spectinomycin or gentamycin for *In-& Out-Cloning*, respectively). Level 0 parts were identified by colony PCR followed by plasmid extraction and subsequent validation by external Sanger sequencing services or by in-house barcoded amplicon-based Nanopore sequencing procedures (13,14). We previously published a stepwise protocol for the basic part generation procedure (17).

**Figure 1.**
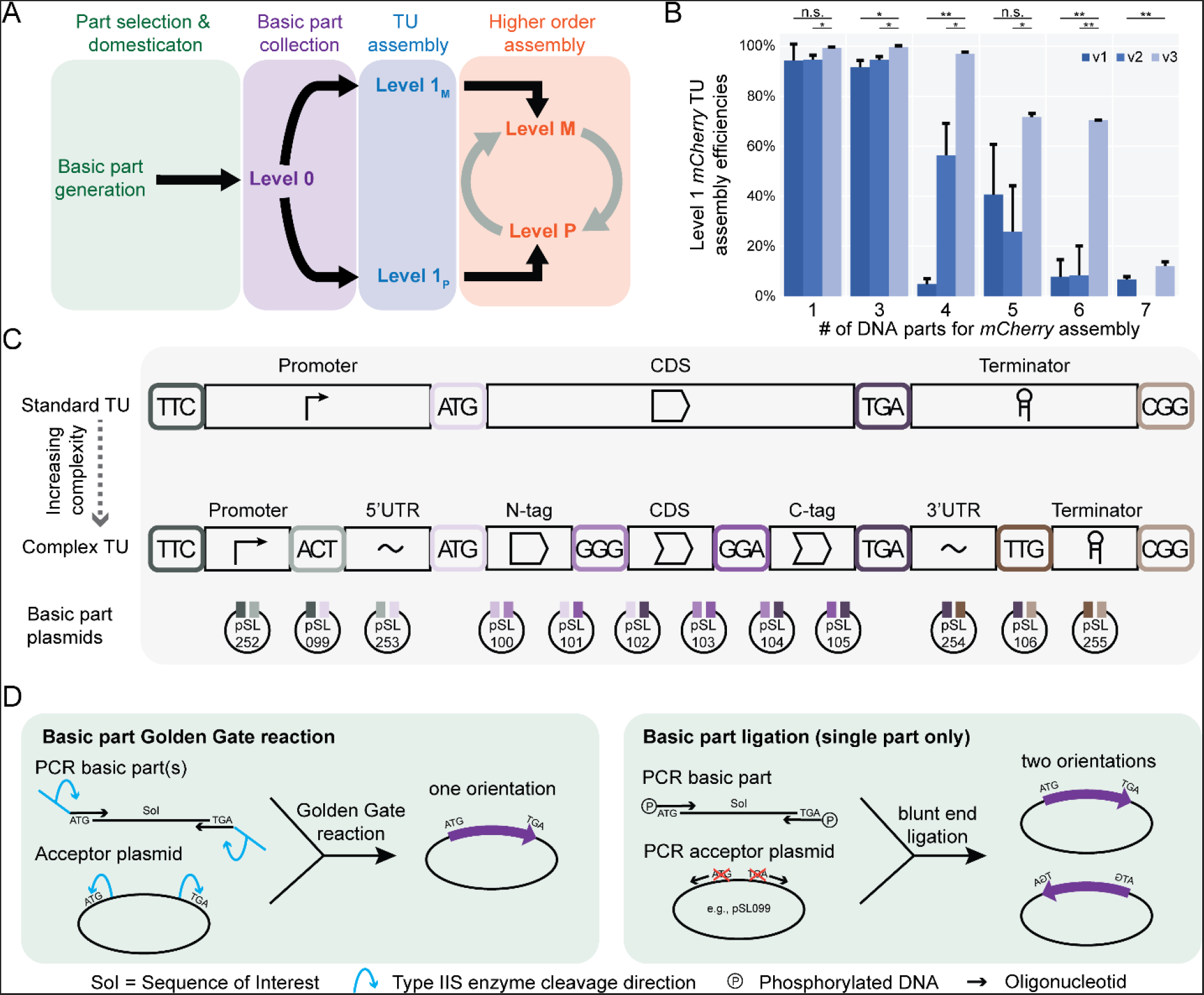
*In-Cloning*: concept, generation of basic parts and transcription units for modular Golden Gate cloning. **(A)** Overview of different levels and the corresponding purpose within modular Golden Gate assemblies. **(B)** Determination of 3 nt fusion site fidelity for the *In-Cloning* workflow using 1 µL total volume Golden Gate reaction mixtures (details see Materials and Methods). Experiments were performed in biological triplicate; on average, 93 colonies per replicate were obtained, the number of colonies obtained decreased with increasing Golden Gate assembly complexity, for CFU data see Table S5. Student’s *t*-test was applied to compare the selected standard (v3) with the standards with lower efficiencies (v1 and v2) (* = *P* < 0.05, ** = *P* < 0.01, n.s. = not significant). **(C)** Overview of Level 0 cloning hierarchy and corresponding plasmid IDs for SapI-based cloning of TUs. Figure S2 shows a more detailed representation of the fusion sites, corresponding parts and Level 0 plasmids. **(D)** Cloning of Level 0 parts can be achieved either by a one pot Golden Gate assembly using BbsI or simply via blunt-end ligation in case of a single phosphorylated DNA fragment.

### Linker design

Linker sequences were designed using SiteOut (18) with the following settings: 1. Sequence Design - Spacer Designer: Spacer_design.txt; 2. Motifs to avoid & GC content: Zipfile with FMs: yeast_pwms.zip, Background GC content: 35%, P value: 0.003, Explicit motifs: Avoid_Motifs.fasta, and GC content: 35%. The respective files are provided in Supporting Data S3. Sequences were ordered as oligonucleotides and plasmids were constructed by Golden Gate DNA assembly resulting in plasmids pSL225-233 and pSL600, details are provided in Table S2, Supporting Data S1 and S2.

### Golden Gate DNA assembly

Golden Gate cloning reactions were performed either in 1 µL total volume using an acoustic dispenser (Echo525 or Echo 650T [Labcyte, San José, USA]) or manually in 10-20 µL total volume. 1 µL reaction mixtures containing 5 fmol of each DNA part, 0.1 μL of the respective Type IIS enzyme (NEB), and 0.1 μL of the T4 ligase (NEB, M0202) in 1× T4 ligase buffer were incubated in a 384-well PCR cycler (Applied Biosystems, Waltham, USA) using the following program: 20 cycles of 4 min at 16 °C, 3 min at 37 °C followed by 10 min at 50 °C, 10 min at 80 °C, and storage at 4 °C. Golden Gate assemblies were then transformed in in-house prepared RbCl competent *E. coli* Top10 cells and 12 transformations were plated by gravity flow of transformation mixture drops on a single 120×120 mm square plate using a 12-channel multi-pipette (12,19). Manual reactions were performed using the same reaction and selection conditions, with the following adjustments to the reaction mix: 50-100 fmol DNA, 1 µL T4 ligase and 1 µL appropriate Type IIS enzyme in 1× T4 ligase buffer in a total volume of 10 to 20 µL. For the construction of basic parts, the following Type IIS enzyme recognition sites are excluded from the sequences: BbsI (BpiI), BsaI, Esp3I (BsmBI) and SapI. We provide stepwise protocols in Köbel and Schindler 2023 and de Vries *et al*. 2024 (17,19).

### Fluorescence protein assay to test Golden Gate assembly fidelity

The coding sequence of *mCherry* was split into varying numbers of parts to determine the SapI-based Golden Gate assembly fidelity in regard to the respective overhangs (for details see Figure S1). Parts were subcloned by blunt-end cloning into Level 0 plasmids resulting in plasmids pSL200-pSL210, and pSL638-pSL650. All subcloned fragments were validated using external Sanger sequencing services. The acceptor plasmid was amplified with appropriate primer pairs (Table S4) providing SapI recognition sites and the designed fusion sites. The amplified product was DpnI digested to remove remaining PCR template prior to BOMB PCR clean-up and subsequent Golden Gate reaction. Golden Gate assemblies were performed as triplicate in 1 µL total volume and transformed into 10 µL chemically competent *E. coli* Top10 cells as described above. Transformation plates were documented using the PhenoBooth (Singer Instruments, UK) using standard settings: Power = 0.5, brightness = 0, gain = 0.01, exposure = 4 ms, hue = 0.5, saturation = 0.19, white balance: Blue = 0.5, red = 0.32, green = 0.22. Assembly fidelity was determined by CFU counting of red and white colonies after transformation of the Golden Gate reactions, the resulting CFU data is available in Table S5.

### Yeast transformation

Yeast strains were transformed according to a modified protocol described by Gietz and Woods (2002) (20). Briefly, yeast strains were inoculated into 5 mL liquid media and incubated overnight at 30 °C with rotation. Overnight cultures were then reinoculated into 20 mL liquid media to OD_600_ = 0.1 and incubated with rotation to a target OD_600_ between 0.5 and 1. Cells were then centrifuged at 2,000 *g* for 5 min, washed with 10 mL sterile ddH_2_O and centrifuged again. Cell pellets were washed with 10 mL 0.1 M LiOAc and centrifuged at 2,000 *g* for 5 min. The cell pellet was resuspended in 400 µL 0.1 M LiOAc. 50 μL cells were mixed with 5 to 10 μL transforming DNA, 36 μL 1M LiOAc, 14 to 19 μL ddH_2_O (depending on DNA volume: DNA + ddH_2_O = 24 µL), 25 μL 10 mg/mL salmon sperm carrier DNA and 240 μL 44% PEG 3350. Transformation mixtures were briefly mixed by inversion and incubated at 30 °C for 30 min before the addition of 36 μL DMSO, followed by inversion and heat shock at 42 °C for 15 min. Cells were pelleted at 1,000 *g* for 1 min, resuspended in 400 μL of 5 mM CaCl_2_, incubated at room temperature for at least 10 min, and plated onto selective media.

### Spotting assay for qualitative β-carotene production test

Yeast strains were streaked onto selective growth plates. Three single colonies of each strain were inoculated into selective media and grown overnight. 200 µL of culture normalized to OD_600_ = 10 were transferred to 96-well flat bottom microtiter plates. The normalized culture was then used to perform a 10-fold dilution series. The Rotor HDA+ (Singer Instruments, UK) was used to pin the dilution series in a 7×7 grid on selective media on a single-well microtiter plate, revisiting the source plate after each pinning step. Incubation of plates prior documentation was performed 48 h at 30 °C, 24 h at room temperature, and 24 h at 6 °C. The experiment was performed in biological triplicate with technical duplicates. Documentation was performed using the PhenoBooth (Singer Instruments, UK) using the following settings: Power = 0.3, brightness = 0, gain = 0, exposure = 4 ms, hue = 0.5, saturation = 0.27, white balance: Blue = 0.35, red = 0.31, green = 0.22.

## Results and Discussion

### In-Cloning: A plasmid toolbox for construction of basic parts and transcription units

The *In-Cloning* plasmid toolbox is designed to generate basic parts in Level 0 plasmids for subsequent assembly into TUs using the enzyme SapI (Figure 1A). SapI has the advantage of a 3 nt fusion site, which may come with a trade-off of reduced cloning fidelity compared to the 4 nt fusion sites generated by commonly used Type IIS enzymes. To identify a set of fusion sites with high fidelity, three different sets of fusion site designs were tested for their assembly fidelity by assembling *mCherry* TUs of varying complexity (Figure 1B, Figure S1), resulting in the depicted overhang hierarchy (Figure 1C, Figure S2). The goal of the plasmid toolbox was to reliably assemble three to five parts which would be either a standard TU consisting of promoter, CDS and terminator or a more complex TU where up to two additional features are added to the assembly (e.g., C- and N-terminal tag). Importantly, the codons ATG and TGA were set by the start and stop codons for the CDS. TGA was chosen because it has stronger Watson-Crick base pairing than TAA, and TAG is often used for the incorporation of non-proteinogenic/non-natural amino acids, for example in the first recoded genome of *E. coli* or in the Synthetic Yeast Project (Sc2.0) (21,22). The fusion sites for fusion proteins (C- and/or N-terminal tagging) were selected as glycine codons. Glycine linkers are naturally occurring and provide flexibility to the fusion protein and are therefore commonly used (23).

The design of the fusion sites was selected based on the ligase fidelity study of Vladimir Potapov and coworkers from 2018 using the fidelity matrices before computational tools were available (5). Three different sets of fusion sites were tested and the one that showed the best fidelity with assembly rates between 75 and 100% for up to 6 basic parts (version 3) was selected to create the plasmid toolbox. This assembly standard fulfills the needs and requirements of the designed assembly hierarchy because at least three out of four candidates are supposed to contain the correct assembled DNA construct. In case of the maximum possible seven part assembly the NEBridge Ligase Fidelity tool (version 1.0) predicts a fidelity of 88% (settings: SapI 37°C/16°C cycling program) for our chosen set. However, in our fidelity test we observed the number of obtained colonies and correct assemblies dropped drastically and do not match the prediction (Figure 1B, Table S5). We traced the drop in assembly fidelity to the TTG fusion site, which may be replaced in the future to optimize the standard. This level of complexity is rarely used and is an option if one is interested in the effects of 3’
s UTRs. Such studies would most likely not use the full complexity of the seven-part assembly. Importantly the assembly fidelity of the selected fusion site hierarchy is sufficient for the intended automated workflows, where normally three (promoter, CDS, terminator) to five parts (promoter, N-terminal tag, CDS, C-terminal tag, terminator) are assembled to TUs. In these scenarios, three out of four candidates should show the correct assembly (Figure 1B).

To reduce background during Golden Gate cloning (caused by remaining acceptor plasmids) the plasmids of the toolbox contains our previously established dual selection cassette expressing *mCherry* and the gyrase inhibitor *ccdB* (12). The expression of *ccdB* causes cell death unless *ccdA* or gyrase suppressor mutations are present in the cloning strain to propagate the acceptor plasmid (e.g., *E. coli* DB3.1). This design has been successfully used previously and allows visual inspection of assemblies in case cells are not sensitive to *ccdB* or mutations impair *ccdB* function (12). Importantly, expression of the *ccdB* toxin enables work with sequence libraries by drastically reducing the background (4,8). An overview of all plasmids and the corresponding plasmid GenBank files can be found in Supporting Data S1 and S2.

Level 0 parts in the plasmid toolbox can be generated with two strategies, either by Golden Gate cloning using BbsI, which allows the combination of multiple sequence fragments (Figure 1D). Alternatively, Level 0 parts can be generated from a single fragment by blunt-end cloning by adding the appropriate fusion sequences to the amplified or synthesized DNA fragment. The respective fragments are phosphorylated and blunt-end ligated to the Level 0 plasmid amplified with universal primers. This strategy results in a forward or reverse orientation within the plasmid. However, this does not affect the subsequent use of the basic parts because the correct fusion sites are attached during the PCR. This approach can speed up and simplify the construction of Level 0 part libraries. To allow modular assembly of TUs, the Level 1 plasmids are fully compatible with the established MoClo system (2) and our previously established MoClo plasmids (8,24,25).

The provided set of Level 1 plasmids also contains the dual selection cassette and allows the combination of Level 0 plasmids to create TUs in forward or reverse orientation (Figure 1A, Figure S3). In addition, the set has been extended to be compatible with Level M and Level P, providing greater flexibility for complex cloning approaches.

### Out-Cloning: An expandable part library and workflow for on demand acceptor plasmid construction

The cloning of Level M and Level P plasmids is intended for the construction of large multigene DNA constructs (2). In theory, they allow for the assembly of DNA fragments of infinite size (Figure 1A and 2A), and we have used them to assemble constructs up to 100,000 bps (25). We have previously generated optimized Level M and P plasmids with their corresponding endlinkers (8,24). These are used in conjunction with our new plasmid toolbox. However, the resulting multigene assemblies in the Level M and P construction plasmids may not be suitable for downstream applications in the targeted organism (24,25). For example, they are not compatible with *Saccharomyces cerevisiae* because they lack the respective autonomously replication sequences (ARS), a centromere and a respective selection marker. This could lead to frequent and labor-intensive construction of acceptor plasmids based on the targeted model organism. To streamline this problem in an economical way, *Out-Cloning* was developed (Figure 2A). *Out-Cloning* is based on a growing collection of plasmid elements that can be combined with flexible linker sequences (or “connectors”) to construct acceptor plasmids quickly and on demand (Figure 2B). The library consists of different positions (e.g. bacterial marker, bacterial origin of replication) which are assembled utilizing flexible linkers allowing to shuffle positions in the acceptor plasmid according to the user’s needs (Figure 2B-C) (note: not all possible linker sequences have been generated, available linkers are listed in Supporting Data S1). The linker set is expanded based on demand and the respective plasmid maps are provide in Supporting Data S2. For the construction of acceptor plasmids, Esp3I (BsmBI isoschizomer), a Type IIS enzyme with 4 nt, was chosen, which is compatible with *In-Cloning* and Level M/P cloning. The basic part Level 0 plasmids are similar to the *In-Cloning* Level 0 plasmids except for the use of the conditional *R6K* origin to allow efficient selection for correctly assembled acceptor plasmids and avoiding background caused by basic part plasmids carrying the resistance marker selected for. The fusion sites were selected based on the aforementioned cloning fidelity tool for optimal cloning results (Figure 2B) (5). An initial basic part library was constructed and is continuously expanded to match application needs (Supporting Data S1 and S2).

**Figure 2.**
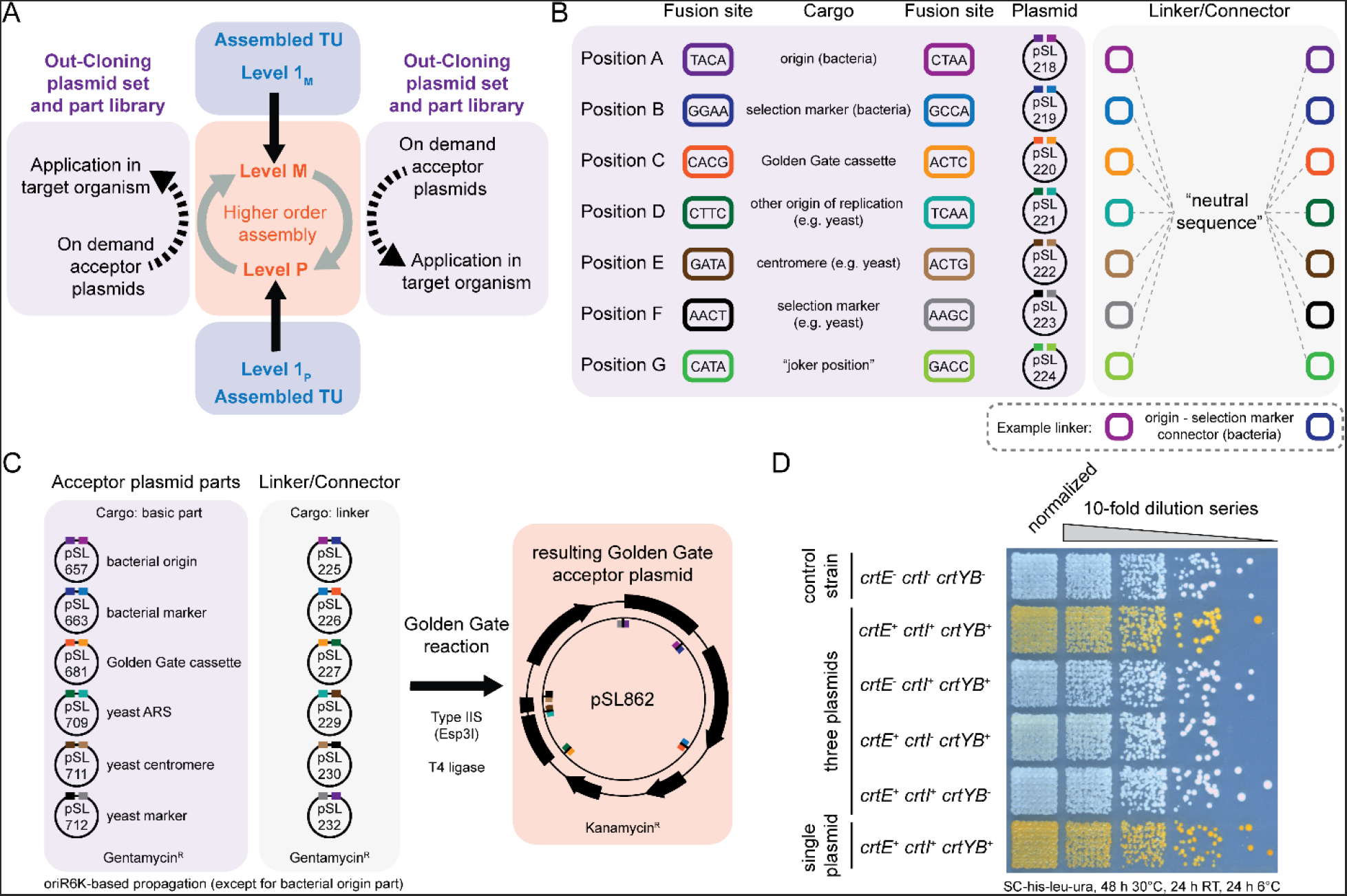
*Out-Cloning*: On demand creation of modular acceptor plasmids and to exit plasmids for modular Golden Gate assembly strategies. **(A)** Level M/P exit-concept cloning strategy. Note: *Out-Cloning* can generate any other type of plasmid (e.g. Level 1 plasmids) based on the user’s needs by extending and combining the basic part libraries. **(B)** From basic elements to modular acceptor plasmids. Linker sequences can be used to combine any position in the forward or reverse direction to generate acceptor plasmids on demand. A set of possible linkers is available (Supporting Data S1 and S2), but not all theoretical possible combinations. **(C)** Workflow of modular acceptor plasmid construction. pSL862 is shown as an example. The colored bars in indicate the overhang sequences for the parts, linkers, and the fusion points based on the linker sequences in the final construct. **(D)** Example application where multiple Golden Gate acceptor plasmids for *S. cerevisiae* were required to test *β*-carotene production on three individual plasmids (pSL885-pSL887) compared to a single plasmid (pSL891). In cases of the indicated absence of a gene, an empty plasmid with a 16 nt neutral spacer sequence (linker sequence of pSL225) was transformed as a control (pSL888-pSL890). Normalization indicates OD_600_ = 10. A representative image is shown. Experiment was performed as biological triplicate with technical duplicate (Figure S4).

*Out-Cloning* provides a rapid framework to construct appropriate Golden Gate acceptor plasmids on demand. Depending on the cloning cassette selected it results in a Level 1, Level M or Level P acceptor plasmid with the additional features selected. As an application example of *Out-Cloning*, four pRS-like (26) yeast Golden Gate acceptor plasmids (three Level 1 [pSL858-pSL860] and one Level P [pSL862]) were assembled. The corresponding workflow is shown in Figure 2C based on the example pSL862. The acceptor plasmids were subsequently used to assemble either the entire *β*-carotene pathway in a single plasmid or each gene in a single plasmid carrying different auxotrophic markers. The rationale behind the tripartite plasmid system would be rapid testing of enzyme variants on TU level instead of combinatorial pathway assemblies.

The plasmid with the complete pathway (pSL891) and the three plasmids with the individual TUs (pSL885-pSL887) were transformed into *S. cerevisiae* BY4741 and *β*-carotene production was observed (Figure 2D, Figure S4). As a control three plasmids containing a 16 nt neutral DNA sequence (linker sequence of pSL225) were assembled (pSL888-pSL890) and combinations of the tripartite plasmid system where one or all of the genes were lacking were generated. As expected, in case of the absence of one or all of the genes no *β*-carotene production was observed. For comparison, the respective empty plasmids were transformed into the strain carrying the entire pathway on a single plasmid allowing for growth on the selective media (SC-his-leu-ura). Only those yeast strains containing the complete pathway, either distributed on one or three plasmids, show pigment formation. Heterogeneity in the β-carotene production of individual colonies was observed for both strains. The heterogeneity appears to be increased in the strain with the pathway on a single plasmid, but the pigment intensity indicates that the overall product accumulation is slightly increased (Figure S4). Nevertheless, the small qualitative differences suggest that a tripartite plasmid system could be used for rapid pathway prototyping or for initial screening purposes prior to assembling complex, larger biosynthetic pathways.

## Conclusion

The *In-& Out-Cloning* plasmid toolboxes are versatile extensions of the MoClo system. *In-Cloning* utilizes the 3 nt fusion sites of the SapI Type IIS enzyme, resulting in minimized fusion sites at the TU level. The provided toolbox allows to build part libraries and allow for TU assembly. Start-Stop cloning has been established previously (3), but a dedicated plasmid toolbox compatible with Modular Cloning implementing new ligase fidelity data has been lacking (5,6). Reduced fusion sites are particularly useful for the cloning and application of non-coding RNA molecules, where proper folding is essential for functionality (4,12,27,28). We have already shown that the 3 nt fusion sites are suitable for generating libraries of synthetic small regulatory RNAs (4). *Out-Coning* allows on-demand construction of acceptor plasmids from a growing library of parts. The provided toolbox allows to expand the part libraries and provides a workflow for the construction acceptor plasmids. This is important when a DNA construct moves from the production step (performed with the standard Level M/P plasmids) to the applied part of a project, which often results in the exchange of the biological model system. Notably, *In-& Out-Cloning* is organism independent and provides the user with a framework for starting part collections beyond the available elements. The system currently conforms to the MoClo standard (2). However, any other standard may be implemented by simply constructing the necessary cloning cassette module(s) as basic parts with the required Type IIS recognition sites and matching fusion sites. The construction of acceptor plasmids based on a characterized library of parts is economical, sustainable, and allows for rapid shuffling of elements when necessary. All plasmids of this study including the previously published Level M, Level P and the corresponding endlinker are available upon request (8,24). Detailed information and annotated sequence data in the form of GenBank files are provided in the Supporting Data S1 and S2.

## Supporting Information

1. Supporting Information – Supplementary Figures, Tables and References (.pdf)
2. Supporting Data S1 – Detailed plasmid information (.xls)
3. Supporting Data S2 – Genbank sequences for all generated plasmids (.zip)
4. Supporting Data S3 – SiteOut files for the generation of neutral linkers (.zip)

## Material Availability

All materials are available from the corresponding author upon request.

## Data Availability

All data are available in the manuscript and its supporting information, or from the corresponding author upon request.

## Acknowledgements

We thank lab members for extensive discussions on building DNA. We thank Torsten Waldminghaus (TU Darmstadt) and Yizhi Cai (University of Manchester) for sharing strains and/or plasmids.

## Author Contributions

DS conceived the study. STdV, TSK, AS and DS conducted the experiments and evaluated all data. STdV and DS wrote the manuscript. All authors read and approved the final manuscript.

## References

1. Engler, C., Kandzia, R. and Marillonnet, S. (2008) A one pot, one step, precision cloning method with high throughput capability. PLoS One, 3, e3647.

2. Weber, E., Engler, C., Gruetzner, R., Werner, S. and Marillonnet, S. (2011) A modular cloning system for standardized assembly of multigene constructs. PLoS One, 6, e16765.

3. Taylor, G.M., Mordaka, P.M. and Heap, J.T. (2019) Start-Stop Assembly: A functionally scarless DNA assembly system optimized for metabolic engineering. Nucleic Acids Res, 47, e17.

4. Brück, M., Köbel, T.S., Dittmar, S., Ramírez Rojas, A.A., Georg, J., Berghoff, B.A. and Schindler, D. (2024) A library-based approach allows systematic and rapid evaluation of seed region length and reveals design rules for synthetic bacterial small RNAs. bioRxiv.

5. Potapov, V., Ong, J.L., Langhorst, B.W., Bilotti, K., Cahoon, D., Canton, B., Knight, T.F., Evans, T.C., Jr. and Lohman, G.J.S. (2018) A single-molecule sequencing assay for the comprehensive profiling of T4 DNA ligase fidelity and bias during DNA end-joining. Nucleic Acids Res, 46, e79.

6. Potapov, V., Ong, J.L., Kucera, R.B., Langhorst, B.W., Bilotti, K., Pryor, J.M., Cantor, E.J., Canton, B., Knight, T.F., Evans, T.C., Jr. et al. (2018) Comprehensive profiling of four base overhang ligation fidelity by T4 DNA ligase and application to DNA assembly. ACS Synth Biol, 7, 2665–2674.

7. Pryor, J.M., Potapov, V., Bilotti, K., Pokhrel, N. and Lohman, G.J.S. (2022) Rapid 40 kb genome construction from 52 Parts through data-optimized assembly design. ACS Synth Biol, 11, 2036–2042.

8. Schindler, D., Milbredt, S., Sperlea, T. and Waldminghaus, T. (2016) Design and assembly of DNA sequence libraries for chromosomal insertion in bacteria based on a set of modified MoClo vectors. ACS Synth Biol, 5, 1362–1368.

9. Kaiser, C., Michaelis, S. and Mitchell, A. (1994) Methods in yeast genetics: A Cold Spring Harbor Laboratory course manual. Cold Spring HarborNY: Cold Spring Harbor Laboratory Press.

10. Zulkower, V. and Rosser, S. (2020) DNA Chisel, a versatile sequence optimizer. Bioinformatics, 36, 4508–4509.

11. Oberacker, P., Stepper, P., Bond, D.M., Hohn, S., Focken, J., Meyer, V., Schelle, L., Sugrue, V.J., Jeunen, G.J., Moser, T. et al. (2019) Bio-On-Magnetic-Beads (BOMB): Open platform for high-throughput nucleic acid extraction and manipulation. PLoS Biol, 17, e3000107.

12. Köbel, T.S., Melo Palhares, R., Fromm, C., Szymanski, W., Angelidou, G., Glatter, T., Georg, J., Berghoff, B.A. and Schindler, D. (2022) An easy-to-use plasmid toolset for efficient generation and benchmarking of synthetic small RNAs in bacteria. ACS Synth Biol, 11, 2989–3003.

13. Ramirez Rojas, A.A., Brinkmann, C.K., Kobel, T.S. and Schindler, D. (2024) DuBA.flow - A low-cost, long-read amplicon sequencing workflow for the validation of synthetic DNA constructs. ACS Synth Biol, 13, 457–465.

14. Ramirez Rojas, A.A., Brinkmann, C.K. and Schindler, D. (2024) Validation of Golden Gate assemblies using highly multiplexed Nanopore amplicon sequencing. arXiv.

15. Gibson, D.G., Young, L., Chuang, R.Y., Venter, J.C., Hutchison, C.A., 3rd and Smith, H.O. (2009) Enzymatic assembly of DNA molecules up to several hundred kilobases. Nat Methods, 6, 343–345.

16. Hanahan, D. (1983) Studies on transformation of Escherichia coli with plasmids. J Mol Biol, 166, 557–580.

17. de Vries, S.T., Kley, L. and Schindler, D. (2024) Use of a Golden Gate plasmid set enabling scarless MoClo-compatible transcription unit assembly. arXiv.

18. Estrada, J., Ruiz-Herrero, T., Scholes, C., Wunderlich, Z. and DePace, A.H. (2016) SiteOut: An online tool to design binding site-free DNA sequences. PLoS One, 11, e0151740.

19. Köbel, T. and Schindler, D. (2023) Automation and miniaturization of Golden Gate DNA assembly reactions using acoustic dispensers. arXiv.

20. Gietz, R.D. and Woods, R.A. (2002) Transformation of yeast by lithium acetate/single-stranded carrier DNA/polyethylene glycol method. Methods Enzymol, 350, 87–96.

21. Isaacs, F.J., Carr, P.A., Wang, H.H., Lajoie, M.J., Sterling, B., Kraal, L., Tolonen, A.C., Gianoulis, T.A., Goodman, D.B., Reppas, N.B. et al. (2011) Precise manipulation of chromosomes in vivo enables genome-wide codon replacement. Science, 333, 348–353.

22. Richardson, S.M., Mitchell, L.A., Stracquadanio, G., Yang, K., Dymond, J.S., DiCarlo, J.E., Lee, D., Huang, C.L., Chandrasegaran, S., Cai, Y. et al. (2017) Design of a synthetic yeast genome. Science, 355, 1040–1044.

23. Reddy Chichili, V.P., Kumar, V. and Sivaraman, J. (2013) Linkers in the structural biology of protein-protein interactions. Protein Sci, 22, 153–167.

24. Messerschmidt, S.J., Schindler, D., Zumkeller, C.M., Kemter, F.S., Schallopp, N. and Waldminghaus, T. (2016) Optimization and characterization of the synthetic secondary chromosome synVicII in Escherichia coli. Front Bioeng Biotechnol, 4, 96.

25. Zumkeller, C., Schindler, D. and Waldminghaus, T. (2018) Modular assembly of synthetic secondary chromosomes. Methods Mol Biol, 1837, 71–94.

26. Sikorski, R.S. and Hieter, P. (1989) A system of shuttle vectors and yeast host strains designed for efficient manipulation of DNA in Saccharomyces cerevisiae. Genetics, 122, 19–27.

27. Philipp, N., Brinkmann, C.K., Georg, J., Schindler, D. and Berghoff, B.A. (2023) DIGGER-Bac: Prediction of seed regions for high-fidelity construction of synthetic small RNAs in bacteria. Bioinformatics, 39.

28. Brück, M., Berghoff, B.A. and Schindler, D. (2024) In silico design, in vitro construction, and in vivo application of synthetic small regulatory RNAs in bacteria. Methods Mol Biol, 2760, 479–507.

