## Supporting Information for "*In- & Out-Cloning*: Plasmid toolboxes for scarless transcription unit and modular Golden Gate acceptor plasmid assembly"

**This file contains:**

Supporting Figures S1 – S4

Supporting Tables S1 – S5

Supporting References

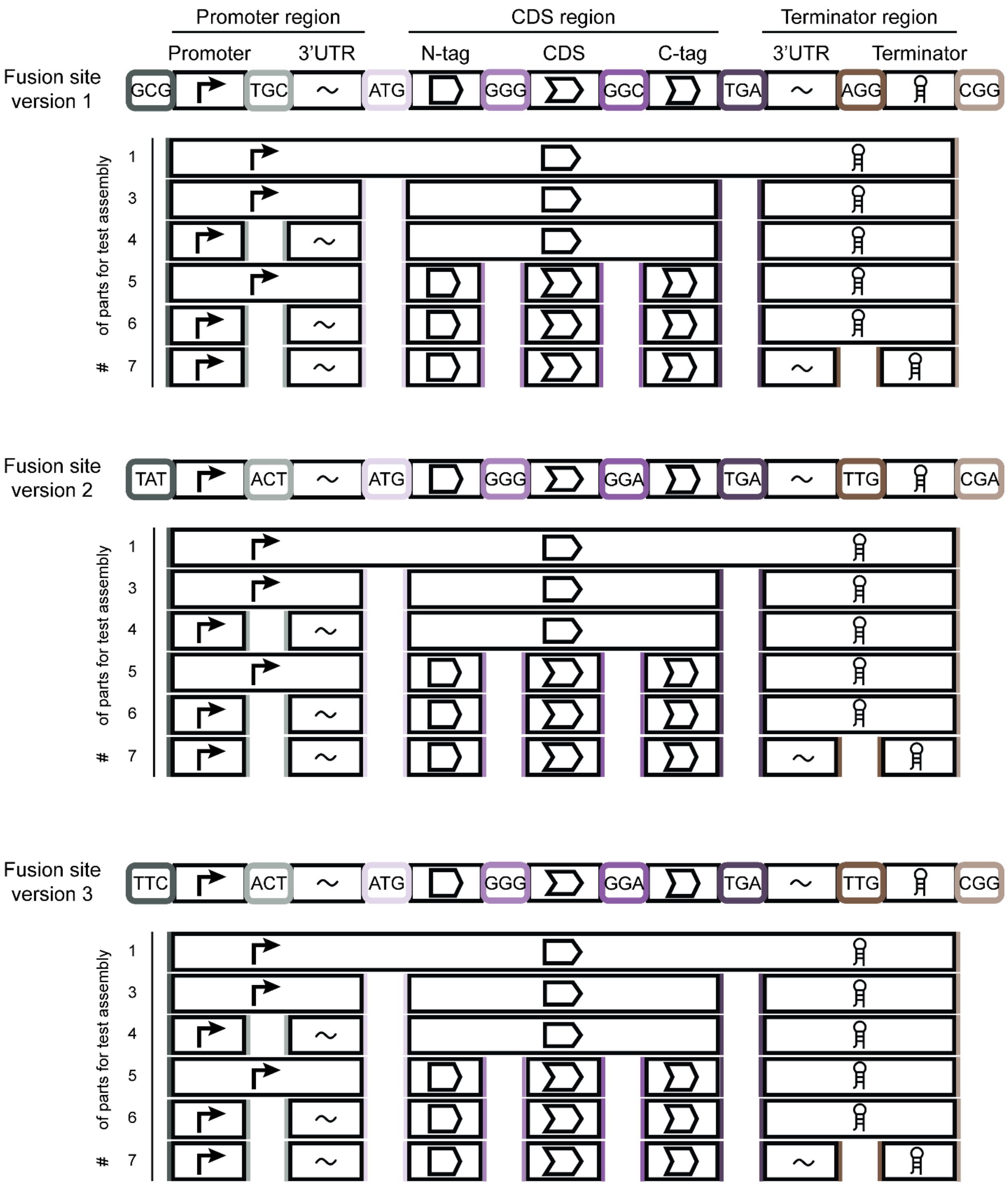

**Figure S1 | Design and fragmentation of *mCherry* transcription unit for the Golden Gate assembly fidelity assessment for 3 nt fusion sites.** Fusion site combinations and the number of parts for each assembly corresponding to Figure 1 B are visualized. CFU data of fidelity assessment is provided in Table S5.

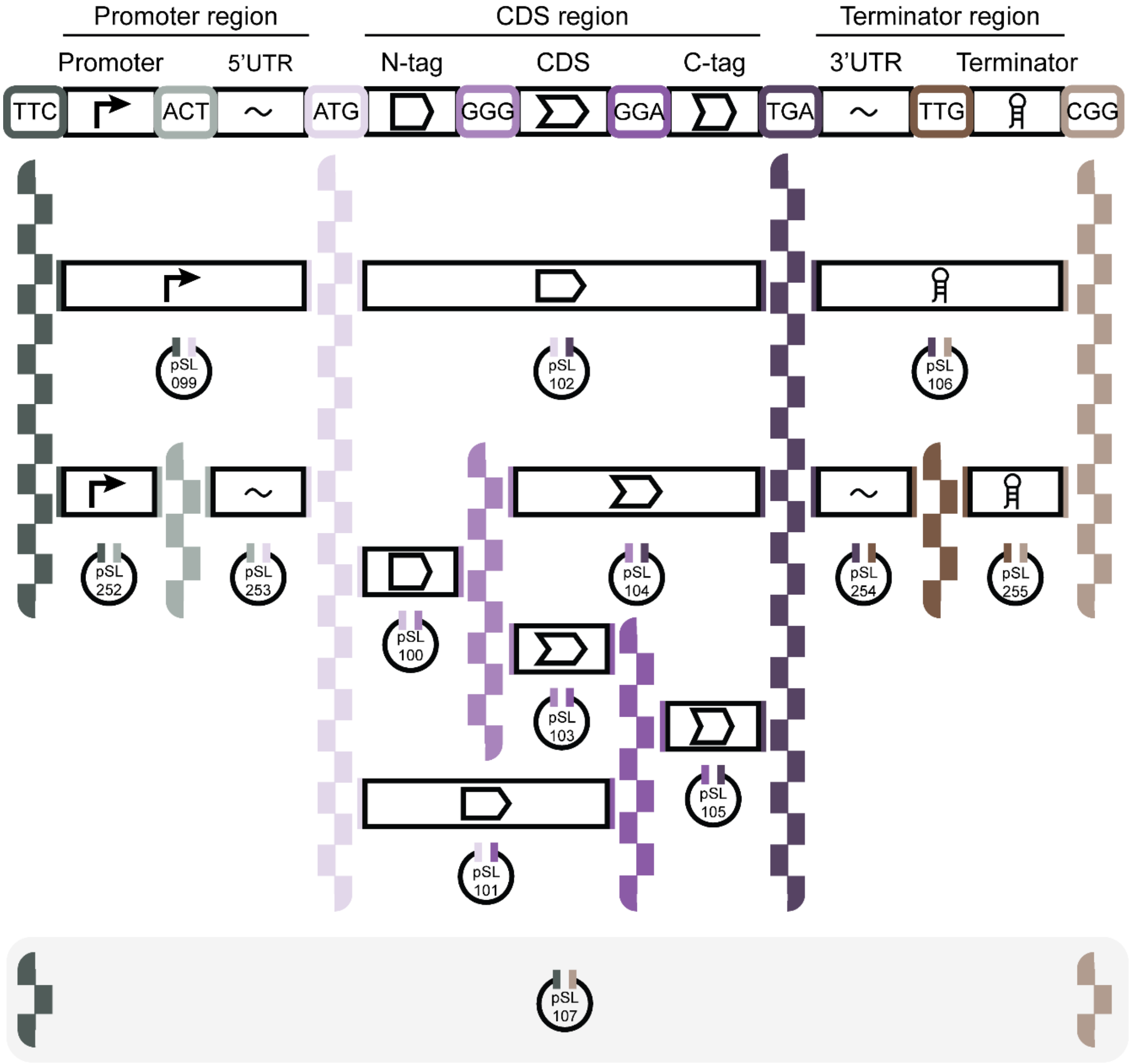

**Figure S2 | Overview of the basic part and acceptor plasmid hierarchy for *In-Cloning* transcription unit generation.** Detailed representation of Figure 1C showing the designed transcription unit hierarchy and its corresponding plasmids. pSL107 is intended to serve as acceptor plasmid for spacer sequences. The color-coded lines between the elements and on the plasmids visualize the fusion site hierarchy. Details of all plasmids can be found in Supporting Data S1 and S2.

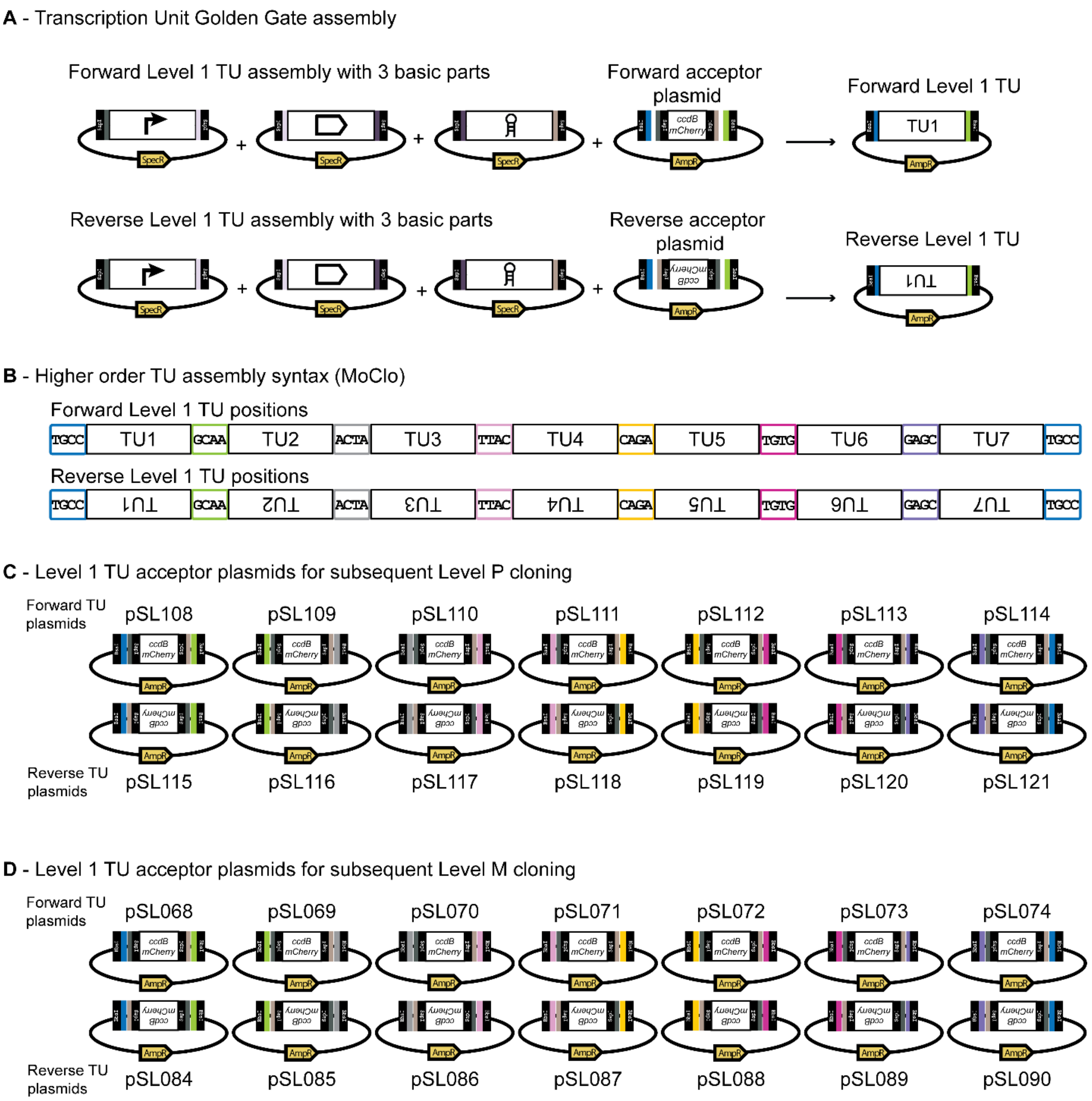

**Figure S3 | Overview of Level 1 TU assembly and respective plasmid sets. (A)** Exemplary assembly of three basic parts of the *In-Cloning* plasmid set in forward (top panel) and reverse (bottom panel) Level 1 acceptor plasmid. Only the orientation of the parts changes not the fusion sites for the higher order assembly in Level M/P. The Level M/P concept and the required endlinker are described in Weber *et al*. 2011, Schindler *et al*. 2016, and Messerschmidt *et al*. 2016 (1-3). All relevant plasmid information can be found in Supporting Data S1 and S2 including the Level M/P and endlinker plasmids. **(B)** Overview of the fusion sites of the used Golden Gate assembly standard established by the group of Sylvestre Marillonnet for higher order assemblies (1). Based on the acceptor plasmid, the TU has a forward or reverse orientation in higher order assemblies. Forward and reverse orientation can be freely mixed and matched. **(C)** Overview of the generated Level 1 acceptor plasmids for higher order assembly in Level P. **(D)** Overview of the generated Level 1 acceptor plasmids for higher order assembly in Level M. Details of all plasmids can be found in Supporting Data S1 and S2.

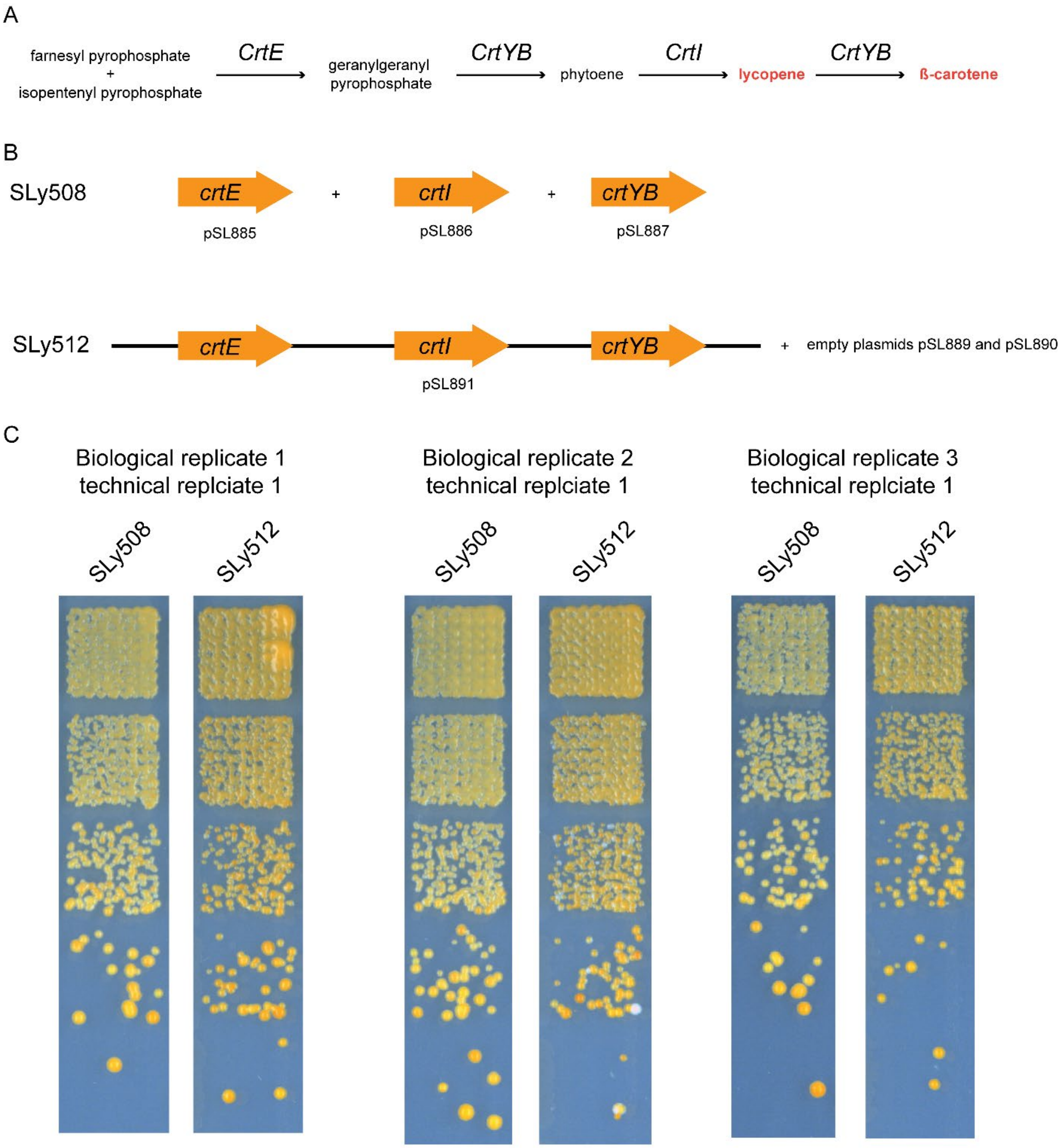

**Figure S4 | Overview of the *β*-carotene pathway and producer strains. (A)** Biosynthetic pathway from farnesyl pyrophosphate and isopentenyl pyrophosphate to *β*-carotene by *CrtE*, *CrtI* and *CrtYB*. Lycopene and *β*-carotene are visible as orange color (indicated by orange letters of compounds). **(B)** *β*-carotene producing strains SLy508 and SLy512 contain the same pathway either on three plasmids (pSL885 to pSL887) or on a single plasmid (pSL891). To compare the performance of the strains under the same conditions, SLy512 has two empty plasmids (pSL889 and pSL890). **(C)** Images of spotted producer cultures, three biological replicates are shown, always the first technical replicate is shown. In general, SLy512 shows more intense pigment accumulation. However, heterogeneity seems to be increased in this strain. Strains from the same replicate were grown on the same plate, the control strains in between have been cropped for better comparison (*cf*. Figure 2D).

**Table S1 | Strains used in this study.**

| **Name** | **Relevant features** | **Reference** |
|---|---|---|
| *E. coli ccd*B Survival 2 $T1^R$ | $F^-$*mcr*A Δ(*mrr-hsd*RMS-*mcr*BC) φ80*lac*ZΔM15 Δ*lac*X74 *rec*A1 *ara*Δ139 Δ(*ara-leu*)7697 *gal*U *gal*K *rps*L($Str^R$) *end*A1 *nup*G *fhu*A::*IS2* | Invitrogen |
| *E. coli* DB3.1 | $F^-$ *gyrA462 endA1 glnV44* Δ(*sr1-recA*) *mcrB mrr hsdS20*($r_B^-$, $m_B^-$) *ara14 galK2 lacY1 proA2 rpsL20*($Str^R$) *xyl5* Δ*leu mtl1* | Invitrogen |
| *E. coli* DB3.1λpir | DB3.1, λ+ | (4) |
| *E. coli* DH5αλpir | $F^-$ Φ80*lacZ*ΔM15 Δ(*lacZYA-argF*) U169 *recA1 endA1 hsdR17* (rK-, mK+) *phoA supE44 thi-1 gyrA96 relA1* $λ^+$ | (5) |
| *E. coli* Top10 | $F^-$ *mcrA* Δ(*mrr-hsdRMS-mcrBC*) φ80*lacZ*ΔM15 Δ*lacX74 nupG recA1 araD139* Δ(*ara-leu*)7697 *galE15 galK16 rpsL*($Str^R$) *endA1* $λ^-$ | Invitrogen |
| *S. cerevisiae* BY4741 | MAT**a** *leu2Δ0 met15Δ0 ura3Δ0 his3Δ1* | (6) |
| *S. cerevisiae* SLy0507 | BY4741 pSL888 pSL889 pSL890 | this study |
| *S. cerevisiae* SLy0508 | BY4741 pSL885 pSL886 pSL887 | this study |
| *S. cerevisiae* SLy0509 | BY4741 pSL888 pSL886 pSL887 | this study |
| *S. cerevisiae* SLy0510 | BY4741 pSL885 pSL889 pSL887 | this study |
| *S. cerevisiae* SLy0511 | BY4741 pSL885 pSL886 pSL890 | this study |
| *S. cerevisiae* SLy0512 | BY4741 pSL891 pSL889 pSL890 | this study |

**Table S2 | Plasmids used in this study (all Golden Gate cloning relevant plasmid files are available as .gbk in the Supporting Data S2).**

| ID | Relevant features | Reference |
|---|---|---|
| pMA34 | Level 1 TU construct acceptor plasmid position 1 | (2) |
| pMA60 | Level M multigene construct acceptor plasmid position 1 | (2) |
| pMA61 | Level M multigene construct acceptor plasmid position 2 | (2) |
| pMA62 | Level M multigene construct acceptor plasmid position 3 | (2) |
| pMA63 | Level M multigene construct acceptor plasmid position 4 | (2) |
| pMA64 | Level M multigene construct acceptor plasmid position 5 | (2) |
| pMA65 | Level M multigene construct acceptor plasmid position 6 | (2) |
| pMA66 | Level M multigene construct acceptor plasmid position 7 | (2) |
| pMA67 | Level P multigene construct acceptor plasmid position 1 | (2) |
| pMA68 | Level P multigene construct acceptor plasmid position 2 | (2) |
| pMA69 | Level P multigene construct acceptor plasmid position 3 | (2) |
| pMA70 | Level P multigene construct acceptor plasmid position 4 | (2) |
| pMA71 | Level P multigene construct acceptor plasmid position 5 | (2) |
| pMA72 | Level P multigene construct acceptor plasmid position 6 | (2) |
| pMA73 | Level P multigene construct acceptor plasmid position 7 | (2) |
| pMA667 | Level M multigene construct endlinker position 1 | (3) |
| pMA668 | Level M multigene construct endlinker position 2 | (3) |
| pMA669 | Level M multigene construct endlinker position 3 | (3) |
| pMA670 | Level M multigene construct endlinker position 4 | (3) |
| pMA671 | Level M multigene construct endlinker position 5 | (3) |
| pMA672 | Level M multigene construct endlinker position 6 | (3) |
| pMA673 | Level M multigene construct endlinker position 7 | (3) |
| pMA674 | Level P multigene construct endlinker position 1 | (3) |
| pMA675 | Level P multigene construct endlinker position 2 | (3) |
| pMA676 | Level P multigene construct endlinker position 3 | (3) |
| pMA677 | Level P multigene construct endlinker position 4 | (3) |
| pMA678 | Level P multigene construct endlinker position 5 | (3) |
| pMA679 | Level P multigene construct endlinker position 6 | (3) |

| ID | Relevant features | Reference |
|---|---|---|
| pMA680 | Level P multigene construct endlinker position 7 | (3) |
| pSL001 | Golden Gate cloning plasmid containing dual selection cassette | (7) |
| pSL099 | Level 0 cloning basic part plasmid; Promoter + 5´UTR (TTC - ATG) | (8) |

**Table S3 | Plasmids created in this study (all plasmid files are available as .gbk in the Supporting Data S2).**

| ID | Relevant features * | Parental plasmid | Reference |
|---|---|---|---|
| pSL068 | Level 1 TU assembly acceptor plasmid position 1; forward orientation for Level M assembly | pMA34/pSL001 | This study |
| pSL069 | Level 1 TU assembly acceptor plasmid position 2; forward orientation for Level M assembly | pMA34/pSL001 | This study |
| pSL070 | Level 1 TU assembly acceptor plasmid position 3; forward orientation for Level M assembly | pMA34/pSL001 | This study |
| pSL071 | Level 1 TU assembly acceptor plasmid position 4; forward orientation for Level M assembly | pMA34/pSL001 | This study |
| pSL072 | Level 1 TU assembly acceptor plasmid position 5; forward orientation for Level M assembly | pMA34/pSL001 | This study |
| pSL073 | Level 1 TU assembly acceptor plasmid position 6; forward orientation for Level M assembly | pMA34/pSL001 | This study |
| pSL074 | Level 1 TU assembly acceptor plasmid position 7; forward orientation for Level M assembly | pMA34/pSL001 | This study |
| pSL084 | Level 1 TU assembly acceptor plasmid position 1; reverse orientation for Level M assembly | pMA34/pSL001 | This study |
| pSL085 | Level 1 TU assembly acceptor plasmid position 2; reverse orientation for Level M assembly | pMA34/pSL001 | This study |
| pSL086 | Level 1 TU assembly acceptor plasmid position 3; reverse orientation for Level M assembly | pMA34/pSL001 | This study |
| pSL087 | Level 1 TU assembly acceptor plasmid position 4; reverse orientation for Level M assembly | pMA34/pSL001 | This study |
| pSL088 | Level 1 TU assembly acceptor plasmid position 5; reverse orientation for Level M assembly | pMA34/pSL001 | This study |
| pSL089 | Level 1 TU assembly acceptor plasmid position 6; reverse orientation for Level M assembly | pMA34/pSL001 | This study |
| pSL090 | Level 1 TU assembly acceptor plasmid position 7; reverse orientation for Level P assembly | pMA34/pSL001 | This study |
| pSL100 | Level 0 cloning basic part plasmid; N-terminal tag (ATG - GGG) | pMA60/pSL001 | This study |
| pSL101 | Level 0 cloning basic part plasmid; CDS for C-terminal tagging (ATG - GGA) | pMA60/pSL001 | This study |
| pSL102 | Level 0 cloning basic part plasmid; CDS (ATG - TGA) | pMA60/pSL001 | This study |
| pSL103 | Level 0 cloning basic part plasmid; CDS for N- and C-terminal tagging (GGG - GGA) | pMA60/pSL001 | This study |
| pSL104 | Level 0 cloning basic part plasmid; CDS for N-terminal tagging (GGG - TGA) | pMA60/pSL001 | This study |

| ID | Relevant features * | Parental plasmid | Reference |
|---|---|---|---|
| pSL105 | Level 0 cloning basic part plasmid; C-terminal tag (GGG - TGA) | pMA60/pSL001 | This study |
| pSL106 | Level 0 cloning basic part plasmid; 3´UTR + terminator (TGA - CGG) | pMA60/pSL001 | This study |
| pSL107 | Level 0 cloning basic part plasmid; single fragment TU (TTC - CGG) | pMA60/pSL001 | This study |
| pSL108 | Level 1 TU assembly acceptor plasmid position 1; forward orientation for Level P assembly | pMA34/pSL001 | This study |
| pSL109 | Level 1 TU assembly acceptor plasmid position 2; forward orientation for Level P assembly | pMA34/pSL001 | This study |
| pSL110 | Level 1 TU assembly acceptor plasmid position 3; forward orientation for Level P assembly | pMA34/pSL001 | This study |
| pSL111 | Level 1 TU assembly acceptor plasmid position 4; forward orientation for Level P assembly | pMA34/pSL001 | This study |
| pSL112 | Level 1 TU assembly acceptor plasmid position 5; forward orientation for Level P assembly | pMA34/pSL001 | This study |
| pSL113 | Level 1 TU assembly acceptor plasmid position 6; forward orientation for Level P assembly | pMA34/pSL001 | This study |
| pSL114 | Level 1 TU assembly acceptor plasmid position 7; forward orientation for Level P assembly | pMA34/pSL001 | This study |
| pSL115 | Level 1 TU assembly acceptor plasmid position 1; reverse orientation for Level P assembly | pMA34/pSL001 | This study |
| pSL116 | Level 1 TU assembly acceptor plasmid position 2; reverse orientation for Level P assembly | pMA34/pSL001 | This study |
| pSL117 | Level 1 TU assembly acceptor plasmid position 3; reverse orientation for Level P assembly | pMA34/pSL001 | This study |
| pSL118 | Level 1 TU assembly acceptor plasmid position 4; reverse orientation for Level P assembly | pMA34/pSL001 | This study |
| pSL119 | Level 1 TU assembly acceptor plasmid position 5; reverse orientation for Level P assembly | pMA34/pSL001 | This study |
| pSL120 | Level 1 TU assembly acceptor plasmid position 6; reverse orientation for Level P assembly | pMA34/pSL001 | This study |
| pSL121 | Level 1 TU assembly acceptor plasmid position 7; reverse orientation for Level P assembly | pMA34/pSL001 | This study |
| pSL200 | Level 0 cloning *mCherry* test assembly part; v1_pLAC (GCG - TGC) | pSL099 | This study |
| pSL201 | Level 0 cloning *mCherry* test assembly part; v1_RBS (TGC - ATG) | pSL099 | This study |
| pSL202 | Level 0 cloning *mCherry* test assembly part; v1_pLAC+RBS (GCG - ATG) | pSL099 | This study |
| pSL203 | Level 0 cloning *mCherry* test assembly part; v1-v2-v3_*mCherry* CDS part A (ATG - GGG) | pSL099 | This study |

| ID | Relevant features * | Parental plasmid | Reference |
|---|---|---|---|
| pSL204 | Level 0 cloning *mCherry* test assembly part; v1_*mCherry* CDS part B (GGG - GGC) | pSL099 | This study |
| pSL205 | Level 0 cloning *mCherry* test assembly part; v1_*mCherry* CDS part C (GGC - TGA) | pSL099 | This study |
| pSL206 | Level 0 cloning *mCherry* test assembly part; v1-v2-v3_*mCherry* CDS (ATG - TGA) | pSL099 | This study |
| pSL207 | Level 0 cloning *mCherry* test assembly part; v1_5'UTR (TGA - AGG) | pSL099 | This study |
| pSL208 | Level 0 cloning *mCherry* test assembly part; v1_Terminator (AGG - CGG) | pSL099 | This study |
| pSL209 | Level 0 cloning *mCherry* test assembly part; v1-v3_5'UTR+Terminator (TGA - CGG) | pSL099 | This study |
| pSL210 | Level 0 cloning *mCherry* test assembly part; v1_Operon (GCG - CGG) | pSL099 | This study |
| pSL218 | Out-Cloning Level 0 basic part acceptor plasmid position 1; bacterial origin (TACA - CTAA) | NA | This study |
| pSL219 | Out-Cloning Level 0 basic part acceptor plasmid position 2; bacterial resistance marker (GGAA - GCCA) | NA | This study |
| pSL220 | Out-Cloning Level 0 basic part acceptor plasmid position 3; MoClo cloning cassette (CACG - ACTC) | NA | This study |
| pSL221 | Out-Cloning Level 0 basic part acceptor plasmid position 4; non-bacterial origin (CTTC - TCAA) | NA | This study |
| pSL222 | Out-Cloning Level 0 basic part acceptor plasmid position 5; centromere (GATA - ACTG) | NA | This study |
| pSL223 | Out-Cloning Level 0 basic part acceptor plasmid position 6; auxotrophic/resistance marker (e.g. for yeast) (AACT - AAGC) | NA | This study |
| pSL224 | Out-Cloning Level 0 basic part acceptor plasmid position 7; "joker position" (CATA - GACC) | NA | This study |
| pSL225 | Out-Cloning Level 0 linker to connect position 1 and 2 (CTAA - GGAA) | NA | This study |
| pSL226 | Out-Cloning Level 0 linker to connect position 2 and 3 (GCCA - CACG) | NA | This study |
| pSL227 | Out-Cloning Level 0 linker to connect position 3 and 4 (ACTC - CTTC) | NA | This study |
| pSL228 | Out-Cloning Level 0 linker to connect position 3 and 1 (ACTC - TACA) | NA | This study |
| pSL229 | Out-Cloning Level 0 linker to connect position 4 and 5 (TCAA - GATA) | NA | This study |
| pSL230 | Out-Cloning Level 0 linker to connect position 5 and 6 (ACTG - AACT) | NA | This study |
| pSL231 | Out-Cloning Level 0 linker to connect position 6 and 7 (AAGC - CATA) | NA | This study |
| pSL232 | Out-Cloning Level 0 linker to connect position 6 and 1 (AAGC - TACA) | NA | This study |
| pSL233 | Out-Cloning Level 0 linker to connect position 7 and 1 (GACC - TACA) | NA | This study |

| ID | Relevant features * | Parental plasmid | Reference |
|---|---|---|---|
| pSL252 | Level 0 cloning basic part acceptor plasmid; promoter (TTC - ACT) | pMA60/pSL001 | This study |
| pSL253 | Level 0 cloning basic part acceptor plasmid; 3´UTR (ACT - ATG) | pMA60/pSL001 | This study |
| pSL254 | Level 0 cloning basic part acceptor plasmid; 5´UTR (TGA - TTG) | pMA60/pSL001 | This study |
| pSL255 | Level 0 cloning basic part acceptor plasmid; terminator (TTG - CGG) | pMA60/pSL001 | This study |
| pSL350 | Level 0 cloning basic part: *rpl18B* promoter (TTC - ATG) | pSL099 | This study |
| pSL415 | Level 0 cloning basic part plasmid: *crtE* CDS (ATG - TGA) | pSL102 | This study |
| pSL416 | Level 0 cloning basic part plasmid: *crtI* CDS (ATG - TGA) | pSL102 | This study |
| pSL417 | Level 0 cloning basic part plasmid: *crtYB* CDS (ATG - TGA) | pSL102 | This study |
| pSL600 | Out-Cloning Level 0 linker to connect position 4 and 6 (TCAA - AACT) | NA | This study |
| pSL638 | Level 0 cloning *mCherry* test assembly part; v2_pLAC (TAT - ACT) | pSL099 | This study |
| pSL639 | Level 0 cloning *mCherry* test assembly part; v2-v3_RBS (ACT - ATG) | pSL099 | This study |
| pSL640 | Level 0 cloning *mCherry* test assembly part; v2_pLAC+RBS (TAT - ATG) | pSL099 | This study |
| pSL641 | Level 0 cloning *mCherry* test assembly part; v2-v3_*mCherry* CDS part B (GGG - GGA) | pSL099 | This study |
| pSL642 | Level 0 cloning *mCherry* test assembly part; v2-v3_*mCherry* CDS part C (GGA - TGA) | pSL099 | This study |
| pSL643 | Level 0 cloning *mCherry* test assembly part; v2-v3_5'UTR (TGA - TTG) | pSL099 | This study |
| pSL645 | Level 0 cloning *mCherry* test assembly part; v2_5'UTR+Terminator (TGA - CGA) | pSL099 | This study |
| pSL646 | Level 0 cloning *mCherry* test assembly part; v2_Operon (TAT - CGA) | pSL099 | This study |
| pSL647 | Level 0 cloning *mCherry* test assembly part; v3_pLAC (TTC - ACT) | pSL099 | This study |
| pSL648 | Level 0 cloning *mCherry* test assembly part; v3_pLAC+RBS (TTC - ATG) | pSL099 | This study |
| pSL649 | Level 0 cloning *mCherry* test assembly part; v3_Terminator (TTG - CGG) | pSL099 | This study |
| pSL650 | Level 0 cloning *mCherry* test assembly part; v3_Operon (TTC - CGG) | pSL099 | This study |
| pSL657 | Out-Cloning Level 0 part position 1: ColE1/pMB1/pUC19 - high copy number (TACA - CTAA) | pSL218 | This study |
| pSL658 | Out-Cloning Level 0 part position 1: pMB1 (pBR3220) - medium copy number (TACA - CTAA) | pSL218 | This study |
| pSL659 | Out-Cloning Level 0 part position 1: pSC101 (TACA - CTAA) | pSL218 | This study |

| ID | Relevant features * | Parental plasmid | Reference |
|---|---|---|---|
| pSL660 | Out-Cloning Level 0 part position 1: p15A (TACA - CTAA) | pSL218 | This study |
| pSL661 | Out-Cloning Level 0 part position 1: F1 origin (TACA - CTAA) | pSL218 | This study |
| pSL662 | Out-Cloning Level 0 part position 2: Spectinomycin resistance marker (GGAA - GCCA) | pSL219 | This study |
| pSL663 | Out-Cloning Level 0 part position 2: Kanamycin resistance marker (GGAA - GCCA) | pSL219 | This study |
| pSL664 | Out-Cloning Level 0 part position 2: Ampicillin resistance marker (GGAA - GCCA) | pSL219 | This study |
| pSL665 | Out-Cloning Level 0 part position 2: Tetracycline resistance marker (GGAA - GCCA) | pSL219 | This study |
| pSL666 | Out-Cloning Level 0 part position 2: Chloramphenicol resistance marker (GGAA - GCCA) | pSL219 | This study |
| pSL667 | Out-Cloning Level 0 part position 3: MoClo cloning cassette Level 1 to Level M release, starting position 1 (CACG - ACTC) | pSL220 | This study |
| pSL668 | Out-Cloning Level 0 part position 3: MoClo cloning cassette Level 1 to Level M release, starting position 2 (CACG - ACTC) | pSL220 | This study |
| pSL669 | Out-Cloning Level 0 part position 3: MoClo cloning cassette Level 1 to Level M release, starting position 3 (CACG - ACTC) | pSL220 | This study |
| pSL670 | Out-Cloning Level 0 part position 3: MoClo cloning cassette Level 1 to Level M release, starting position 4 (CACG - ACTC) | pSL220 | This study |
| pSL671 | Out-Cloning Level 0 part position 3: MoClo cloning cassette Level 1 to Level M release, starting position 5 (CACG - ACTC) | pSL220 | This study |
| pSL672 | Out-Cloning Level 0 part position 3: MoClo cloning cassette Level 1 to Level M release, starting position 6 (CACG - ACTC) | pSL220 | This study |
| pSL673 | Out-Cloning Level 0 part position 3: MoClo cloning cassette Level 1 to Level M release, starting position 7 (CACG - ACTC) | pSL220 | This study |
| pSL674 | Out-Cloning Level 0 part position 3: MoClo cloning cassette Level 1 reverse to Level M release, starting position 1 (CACG - ACTC) | pSL220 | This study |
| pSL675 | Out-Cloning Level 0 part position 3: MoClo cloning cassette Level 1 reverse to Level M release, starting position 2 (CACG - ACTC) | pSL220 | This study |
| pSL676 | Out-Cloning Level 0 part position 3: MoClo cloning cassette Level 1 reverse to Level M release, starting position 3 (CACG - ACTC) | pSL220 | This study |
| pSL677 | Out-Cloning Level 0 part position 3: MoClo cloning cassette Level 1 reverse to Level M release, starting position 4 (CACG - ACTC) | pSL220 | This study |
| pSL678 | Out-Cloning Level 0 part position 3: MoClo cloning cassette Level 1 reverse to Level M release, starting position 5 (CACG - ACTC) | pSL220 | This study |
| pSL679 | Out-Cloning Level 0 part position 3: MoClo cloning cassette Level 1 reverse to Level M release, starting position 6 (CACG - ACTC) | pSL220 | This study |

| ID | Relevant features * | Parental plasmid | Reference |
|---|---|---|---|
| pSL680 | Out-Cloning Level 0 part position 3: MoClo cloning cassette Level 1 reverse to Level M release, starting position 7 (CACG - ACTC) | pSL220 | This study |
| pSL681 | Out-Cloning Level 0 part position 3: MoClo cloning cassette Level 1 to Level P release, starting position 1 (CACG - ACTC) | pSL220 | This study |
| pSL682 | Out-Cloning Level 0 part position 3: MoClo cloning cassette Level 1 to Level P release, starting position 2 (CACG - ACTC) | pSL220 | This study |
| pSL683 | Out-Cloning Level 0 part position 3: MoClo cloning cassette Level 1 to Level P release, starting position 3 (CACG - ACTC) | pSL220 | This study |
| pSL684 | Out-Cloning Level 0 part position 3: MoClo cloning cassette Level 1 to Level P release, starting position 4 (CACG - ACTC) | pSL220 | This study |
| pSL685 | Out-Cloning Level 0 part position 3: MoClo cloning cassette Level 1 to Level P release, starting position 5 (CACG - ACTC) | pSL220 | This study |
| pSL686 | Out-Cloning Level 0 part position 3: MoClo cloning cassette Level 1 to Level P release, starting position 6 (CACG - ACTC) | pSL220 | This study |
| pSL687 | Out-Cloning Level 0 part position 3: MoClo cloning cassette Level 1 to Level P release, starting position 7 (CACG - ACTC) | pSL220 | This study |
| pSL688 | Out-Cloning Level 0 part position 3: MoClo cloning cassette Level 1 reverse to Level P release, starting position 1 (CACG - ACTC) | pSL220 | This study |
| pSL689 | Out-Cloning Level 0 part position 3: MoClo cloning cassette Level 1 reverse to Level P release, starting position 2 (CACG - ACTC) | pSL220 | This study |
| pSL690 | Out-Cloning Level 0 part position 3: MoClo cloning cassette Level 1 reverse to Level P release, starting position 3 (CACG - ACTC) | pSL220 | This study |
| pSL691 | Out-Cloning Level 0 part position 3: MoClo cloning cassette Level 1 reverse to Level P release, starting position 4 (CACG - ACTC) | pSL220 | This study |
| pSL692 | Out-Cloning Level 0 part position 3: MoClo cloning cassette Level 1 reverse to Level P release, starting position 5 (CACG - ACTC) | pSL0220 | This study |
| pSL693 | Out-Cloning Level 0 part position 3: MoClo cloning cassette Level 1 reverse to Level P release, starting position 6 (CACG - ACTC) | pSL220 | This study |
| pSL694 | Out-Cloning Level 0 part position 3: MoClo cloning cassette Level 1 reverse to Level P release, starting position 7 (CACG - ACTC) | pSL220 | This study |
| pSL695 | Out-Cloning Level 0 part position 3: MoClo cloning cassette Level M, starting position 1 (CACG - ACTC) | pSL220 | This study |
| pSL696 | Out-Cloning Level 0 part position 3: MoClo cloning cassette Level M, starting position 2 (CACG - ACTC) | pSL220 | This study |
| pSL697 | Out-Cloning Level 0 part position 3: MoClo cloning cassette Level M, starting position 3 (CACG - ACTC) | pSL220 | This study |
| pSL698 | Out-Cloning Level 0 part position 3: MoClo cloning cassette Level M, starting position 4 (CACG - ACTC) | pSL220 | This study |

| ID | Relevant features * | Parental plasmid | Reference |
|---|---|---|---|
| pSL699 | Out-Cloning Level 0 part position 3: MoClo cloning cassette Level M, starting position 5 (CACG - ACTC) | pSL220 | This study |
| pSL700 | Out-Cloning Level 0 part position 3: MoClo cloning cassette Level M, starting position 6 (CACG - ACTC) | pSL220 | This study |
| pSL701 | Out-Cloning Level 0 part position 3: MoClo cloning cassette Level M, starting position 7 (CACG - ACTC) | pSL220 | This study |
| pSL702 | Out-Cloning Level 0 part position 3: MoClo cloning cassette Level P, starting position 1 (CACG - ACTC) | pSL220 | This study |
| pSL703 | Out-Cloning Level 0 part position 3: MoClo cloning cassette Level P, starting position 2 (CACG - ACTC) | pSL220 | This study |
| pSL704 | Out-Cloning Level 0 part position 3: MoClo cloning cassette Level P, starting position 3 (CACG - ACTC) | pSL220 | This study |
| pSL705 | Out-Cloning Level 0 part position 3: MoClo cloning cassette Level P, starting position 4 (CACG - ACTC) | pSL220 | This study |
| pSL706 | Out-Cloning Level 0 part position 3: MoClo cloning cassette Level P, starting position 5 (CACG - ACTC) | pSL220 | This study |
| pSL707 | Out-Cloning Level 0 part position 3: MoClo cloning cassette Level P, starting position 6 (CACG - ACTC) | pSL220 | This study |
| pSL708 | Out-Cloning Level 0 part position 3: MoClo cloning cassette Level P, starting position 7 (CACG - ACTC) | pSL220 | This study |
| pSL709 | Out-Cloning Level 0 part position 4: *S. cerevisiae ARS4* (CTTC - TCAA) | pSL221 | This study |
| pSL710 | Out-Cloning Level 0 part position 4: *S. cerevisiae* 2micron origin (CTTC - TCAA) | pSL221 | This study |
| pSL711 | Out-Cloning Level 0 part position 5: *S. cerevisiae CEN6* (GATA - ACTG) | pSL222 | This study |
| pSL712 | Out-Cloning Level 0 part position 6: *S. cerevisiae HIS3* (AACT - AAGC) | pSL223 | This study |
| pSL713 | Out-Cloning Level 0 part position 6: *S. cerevisiae URA3* (AACT - AAGC) | pSL223 | This study |
| pSL714 | Out-Cloning Level 0 part position 6: *S. cerevisiae LEU2* (AACT - AAGC) | pSL223 | This study |
| pSL715 | Out-Cloning Level 0 part position 6: *KanMX* (AACT - AAGC) | pSL223 | This study |
| pSL716 | Out-Cloning Level 0 part position 6: Zeocin resistance marker (AACT - AAGC) | pSL223 | This study |
| pSL717 | Out-Cloning Level 0 part position 6: Hygromcycin resistance marker (AACT - AAGC) | pSL223 | This study |
| pSL718 | Out-Cloning Level 0 part position 7: *oriT* (CATA - GACC) | pSL224 | This study |

| ID | Relevant features * | Parental plasmid | Reference |
|---|---|---|---|
| pSL858 | Level 1 TU assembly acceptor plasmid position 1 forward orientation for Level P (TTC-CGG) | NA | This study |
| pSL859 | Level 1 TU assembly acceptor plasmid position 1 forward orientation for Level P (TTC-CGG) | NA | This study |
| pSL860 | Level 1 TU assembly acceptor plasmid position 1 forward orientation for Level P (TTC-CGG) | NA | This study |
| pSL862 | Level P pRSII413-like acceptor plasmid for *S. cerevisiae* position 1 for Level M (TGCC-GGGA) | NA | This study |
| pSL882 | Exit-plasmid Level 1 TU assembly plasmid position 1 forward orientation for Level P: pRPL18B-*crtE*-tADH1 (TGCC - GCAA) | pSL108 | This study |
| pSL883 | Exit-plasmid Level 1 TU assembly plasmid position 2 forward orientation for Level P: pRPL18B-*crtI*-tADH1 (GCAA - ACTA) | pSL109 | This study |
| pSL884 | Exit-plasmid Level 1 TU assembly plasmid position 3 forward orientation for Level P. pRPL18B-*crtYB*-tADH1 (ACTA-TTAC) | pSL110 | This study |
| pSL885 | Exit-plasmid Level 1 TU assembly plasmid position 1 forward orientation for Level P. Contains medium expressed *crtE* (TGCC - GCAA) | pSL858 | This study |
| pSL886 | Exit-plasmid Level 1 TU assembly plasmid position 1 forward orientation for Level P. Contains medium expressed *crtI* (TGCC - GCAA) | pSL859 | This study |
| pSL887 | Exit-plasmid Level 1 TU assembly plasmid position 1 forward orientation for Level P. Contains medium expressed *crtYB* (TGCC - GCAA) | pSL860 | This study |
| pSL888 | Exit-plasmid Level 1 TU assembly plasmid position 1 forward orientation for Level P. Contains neutral DNA fragment (TGCC - GCAA) | pSL858 | This study |
| pSL889 | Exit-plasmid Level 1 TU assembly plasmid position 1 forward orientation for Level P. Contains neutral DNA fragment (TGCC - GCAA) | pSL859 | This study |
| pSL890 | Exit-plasmid Level 1 TU assembly plasmid position 1 forward orientation for Level P. Contains neutral DNA fragment (TGCC - GCAA) | pSL860 | This study |
| pSL891 | Exit-plasmid Level P construct for lycopene production under medium strength promoter (TGCC - TTAC) | pSL862 | This study |
| pSL892 | Level 0 cloning basic part plasmid; *adh1* terminator (TGA - CGG) | pSL106 | This study |

* Detailed description of the created plasmids are provided in the Supporting Data S1 and plasmid sequences are provided as GenBank files in Supporting Data S2.

**Table S4 | Oligonucleotides used in this study.**

| Name | Sequence (5´-3´) | Information |
|---|---|---|
| SLo0017 | ATATGCTCTTCTCGCGGCATTGTCTTCACAGAGTG | Primer to amplify Level 1 plasmid for Golden Gate cloning fidelity test of versions 1 and 3 in combination with oligo SLo0018 and SLo0497, respectively |
| SLo0018 | TATAGCTCTTCACGGGCAATTGTCTTCTGCACG | Primer to amplify Level 1 plasmid for Golden Gate cloning fidelity test of version 1 in combination with oligo SLo0017 |
| SLo0125 | TATAGCTCTTCACGAGCAATTGTCTTCTGCACG | Primer to amplify Level 1 plasmid for Golden Gate cloning fidelity test of version 2 in combination with oligo SLo0126 |
| SLo0126 | ATATGCTCTTCTATAGGCATTGTCTTCACAGAGTG | Primer to amplify Level 1 plasmid for Golden Gate cloning fidelity test of version 2 in combination with oligo SLo0125 |
| SLo0128 | GGTTATTGTCTCATGAGCGG | Universal Level 1 colony PCR primer in combination with SLo2120. Also functions as sequencing oligo. |
| SLo0497 | ATATGCTCTTCTGAAGGCATTGTCTTCACAGAGTG | Primer to amplify Level 1 plasmid for Golden Gate cloning fidelity test of version 3 in combination with oligo SLo0017 |
| SLo0765 | TGAAGAGCAGGCACGAACCC | Primer to amplify Level 0 plasmid for blunt cloning in combination with SLo0766. |
| SLo0766 | AGAAGAGCGAGCACAGAGTGC | Primer to amplify Level 0 plasmid for blunt cloning in combination with SLo0765. |
| SLo1496 | GTTCGGTCAAGGTTCTGGAC | Universal Level 0, Level 1 and Out-Cloning colony PCR primer in combination with SLo2120. Also functions as sequencing oligo. |
| SLo2120 | TGGAAAAACGCCAGCAACGC | Universal Level 1 and Level 0 colony PCR primer in combination with SLo1496. Also functions as sequencing oligo. |

**Table S5 | CFU documentation of *In-Cloning* fusion site fidelity determination for the three fusion site versions; red (correct) and white (incorrect) TU assemblies.**

| Assembly* | | CFU | | | | | | Total CFUs | Fidelity [%] | SD [+/- %] |
|---|---|---|---|---|---|---|---|---|---|---|
| | | Replicate 1 | | Replicate 2 | | Replicate 3 | | | | |
| | | Red | White | Red | White | Red | White | | | |
| Version 1 | 1 part | 136 | 1 | 90 | 16 | 348 | 5 | 596 | 94.3 | 8.1 |
| | 3 parts | 267 | 29 | 645 | 30 | 190 | 23 | 1184 | 91.7 | 3.4 |
| | 4 parts | 3 | 73 | 1 | 38 | 13 | 153 | 281 | 4.8 | 2.7 |
| | 5 parts | 15 | 17 | 8 | 5 | 3 | 19 | 67 | 40.7 | 2.5 |
| | 6 parts | 37 | 175 | 3 | 95 | 2 | 74 | 386 | 7.7 | 8.4 |
| | 7 parts | 7 | 122 | 3 | 44 | 7 | 78 | 261 | 6.7 | 1.4 |
| Version 2 | 1 part | 68 | 2 | 73 | 5 | 52 | 4 | 204 | 94.5 | 2.3 |
| | 3 parts | 49 | 3 | 52 | 4 | 26 | 1 | 135 | 94.5 | 1.7 |
| | 4 parts | 4 | 5 | 5 | 5 | 26 | 9 | 54 | 56.2 | 15.9 |
| | 5 parts | 0 | 11 | 6 | 9 | 6 | 10 | 42 | 25.8 | 22.4 |
| | 6 parts | 0 | 3 | 0 | 4 | 2 | 6 | 15 | 8.3 | 14.4 |
| | 7 parts | 0 | 3 | 0 | 5 | 0 | 6 | 14 | 0.0 | 0.0 |
| Version 3 | 1 part | 215 | 1 | 228 | 3 | 224 | 1 | 672 | 99.3 | 0.5 |
| | 3 parts | 71 | 1 | 50 | 0 | 149 | 0 | 271 | 99.5 | 0.8 |
| | 4 parts | 132 | 3 | 150 | 6 | 197 | 6 | 494 | 97.0 | 0.8 |
| | 5 parts | 55 | 20 | 51 | 20 | 51 | 22 | 219 | 71.7 | 1.7 |
| | 6 parts | 19 | 8 | 24 | 6 | 9 | 3 | 69 | 75.1 | 4.8 |
| | 7 parts | 1 | 9 | 3 | 22 | 3 | 18 | 56 | 12.1 | 2.1 |

* corresponding fusion sites for each assembly are visualized in Figure S1.

### Supporting References

1. Weber, E., Engler, C., Gruetzner, R., Werner, S. and Marillonnet, S. (2011) A modular cloning system for standardized assembly of multigene constructs. *PLoS One*, **6**, e16765.
2. Schindler, D., Milbredt, S., Sperlea, T. and Waldminghaus, T. (2016) Design and assembly of DNA sequence libraries for chromosomal insertion in bacteria based on a set of modified MoClo vectors. *ACS Synth Biol*, **5**, 1362-1368.
3. Messerschmidt, S.J., Schindler, D., Zumkeller, C.M., Kemter, F.S., Schallopp, N. and Waldminghaus, T. (2016) Optimization and characterization of the synthetic secondary chromosome synVicII in *Escherichia coli*. *Front Bioeng Biotechnol*, **4**, 96.
4. House, B.L., Mortimer, M.W. and Kahn, M.L. (2004) New recombination methods for *Sinorhizobium meliloti* genetics. *Appl Environ Microbiol*, **70**, 2806-2815.
5. Miller, V.L. and Mekalanos, J.J. (1988) A novel suicide vector and its use in construction of insertion mutations: Osmoregulation of outer membrane proteins and virulence determinants in *Vibrio cholerae* requires *toxR*. *J Bacteriol*, **170**, 2575-2583.
6. Brachmann, C.B., Davies, A., Cost, G.J., Caputo, E., Li, J., Hieter, P. and Boeke, J.D. (1998) Designer deletion strains derived from *Saccharomyces cerevisiae* S288C: A useful set of strains and plasmids for PCR-mediated gene disruption and other applications. *Yeast*, **14**, 115-132.
7. Köbel, T.S., Melo Palhares, R., Fromm, C., Szymanski, W., Angelidou, G., Glatter, T., Georg, J., Berghoff, B.A. and Schindler, D. (2022) An easy-to-use plasmid toolset for efficient generation and benchmarking of synthetic small RNAs in bacteria. *ACS Synth Biol*, **11**, 2989-3003.
8. Brück, M., Berghoff, B.A. and Schindler, D. (2024) *In silico* design, *in vitro* construction, and *in vivo* application of synthetic small regulatory RNAs in bacteria. *Methods Mol Biol*, **2760**, 479-507.
